# Machine intelligence design of 2019-nCoV drugs

**DOI:** 10.1101/2020.01.30.927889

**Authors:** Kaifu Gao, Duc Duy Nguyen, Rui Wang, Guo-Wei Wei

## Abstract

Wuhan coronavirus, called 2019-nCoV, is a newly emerged virus that infected more than 9692 people and leads to more than 213 fatalities by January 30, 2020. Currently, there is no effective treatment for this epidemic. However, the viral protease of a coronavirus is well-known to be essential for its replication and thus is an effective drug target. Fortunately, the sequence identity of the 2019-nCoV protease and that of severe-acute respiratory syndrome virus (SARS-CoV) is as high as 96.1%. We show that the protease inhibitor binding sites of 2019-nCoV and SARS-CoV are almost identical, which means all potential anti-SARS-CoV chemotherapies are also potential 2019-nCoV drugs. Here, we report a family of potential 2019-nCoV drugs generated by a machine intelligence-based generative network complex (GNC). The potential effectiveness of treating 2019-nCoV by using some existing HIV drugs is also analyzed.

## 1 Introduction

A cluster of pneumonia cases of unknown cause emerged with connections to Huanan South China Seafood Market in Wuhan city, Hubei Province of China in late December 2019. On January 7, 2020, a positive-sense, single-stranded RNA coronavirus (CoV) was identified as the causative agent, and the World Health Organization (WHO) named this novel coronavirus as 2019-nCoV on January 10. By January 30, a total of 9692 confirmed cases with 213 deaths had been reported in the world. As reported in,^1^ in comparison with the basic reproduction number (*R*_0_) of the severe acute respiratory syndrome (SARS) epidemic, which is about 4.91, *R*_0_ of the 2019-nCoV epidemic was estimated as high as 6.47, which implies the severity of this widespread dissemination caused by 2019-nCoV. Under the current health emergency, many researchers around the world engaged in the investigation of the genetic and functional data of 2019-nCoV and compare to other coronaviruses to design proper infection control strategy^2,3^ and seek potential drugs that can prevent and/or cure this serious epidemic.^4–7^

On January 10, the genome sequence of 2019-nCoV was first released on GenBank (accession MN908947) by Yong-Zhen Zhang’s group at Fudan University.^8^ Subsequently, phylogenetic analysis revealed that 2019-nCoV belonged to the Betacoronavirus genera and its closest whole-genome relatives are two SARS-like coronaviruses from bats, i.e., ZC45 and ZXC21, which shared about 89% sequence identity with 2019-nCoV.^6,7^ Moreover, detailed sequence alignment data indicates that there is significant genetics distance between the 2019-nCoV and the human SARS-CoV. Therefore, the 2019-nCoV should be considered as a new type of bat coronavirus. However, this observation raised a worth-thinking issue whether the 2019-nCoV has the same infective mechanisms as the human SARS-CoV.

The spike protein (S-protein), which is known as the multi-functional molecular machine that mediates coronavirus entry into host cells^9^ is discussed to answer the question mentioned above. According to the literature,^5,6^ the 2019-nCoV and the SARS-CoV have a high amino acid sequence homology of the S-protein, which can result in severe human infections. Further structural similarity analysis showed and confirmed that the S-protein of 2019-nCoV uses the same human cell receptor angiotensin-converting enzyme 2 (ACE2) as SARS-CoV.^4,10^ Therefore, similar approaches can be applied to prevent the S-protein binding with the receptor ACE2.

S-protein is one of the four proteins in coronavirus, which can be cleaved by a host cell furin-like protease (non-structural protein in coronavirus) into two functional units, S1 and S2. S1 makes up the large receptor-binding domain (RBD) to facilitates virus infection by binding to host receptors and S2 forms the stalk of the spike molecule.^11^ Therefore, one possible way to control the infection is to seek protease inhibitors to prevent the S-protein of 2019-nCoV cleaved into S1. Another possible way is to seek specific drugs that have the ability to inhibit RBD binding to host receptors. However, consider the fact that the diversity of CoV is reflected in the variable S-proteins and the gene of S-protein is easy to mutate,^12–15^ it is not economical to design a new drug to prevent the RBD binding to host receptor directly. Moreover, considering the high sequence identity between viral proteases of 2019-nCoV and SARS-CoV, seeking protease inhibitors to treat respiratory diseases induced by SARS and this novel pneumonia will be our first choice.

Viral proteases are common targets in dealing with human viruses such as the HIV virus and hepatitis C virus. Protease inhibitors are remarkably effective in blocking the replication of coronavirus, including the SARS and the Middle East respiratory syndrome (MERS), providing a promising foundation for the development of anticoronaviral therapeutics. An overview of SARS-CoV 3CL protease inhibitors has been reported.^16^ However, currently, there are no effective anti-SARS-CoV drugs available despite there are more than 3500 publications. It is true that there were only 8422 infected cases and 916 deaths reported for SARS-CoV, which might make the drug development unprofitable. The fact that we are unprepared for another coronaviral pandemic, like the Wuhan coronavirus outbreak, causes the world economy trillion dollars, not to mention the loss of many lives.

The present work makes use of a recently developed generative network complex (GNC)^17^ to explore potential protease inhibitors for curing 2019-nCoV. We generate anticoronaviral therapeutic candidates for 2019-nCoV and evaluate their druggable properties. We also examine the potential of repurposing HIV protease inhibitors, Aluvia and Norvir, for 2019-nCoV.

## 2 Methods

New anti-CoV drug candidates are designed by using a recently developed generative network complex (GNC) platform.^17^ As shown in Fig. 1, the first component is a generative network including an encoder, a latent space, a molecule generator, and a decoder. The generative network will take a given SMILES string as the input to generate novel molecules in terms of SMILES strings, which will be fed into the second component of our GNC, a two-dimensional (2D) fingerprint-based deep neural network (2DFP-DNN), to reevaluate their druggable properties. The next component is the MathPose model which is used to predict the three-dimensional (3D) structure information of the compounds selected by 2DFP-DNN. The bioactivities of those compounds are further estimated by the structure-based deep learning model named MathDL.^18^ The druggable properties predicted by this last component of our GNC are used as an indicator to select the promising drug candidates.

**Figure 1:**
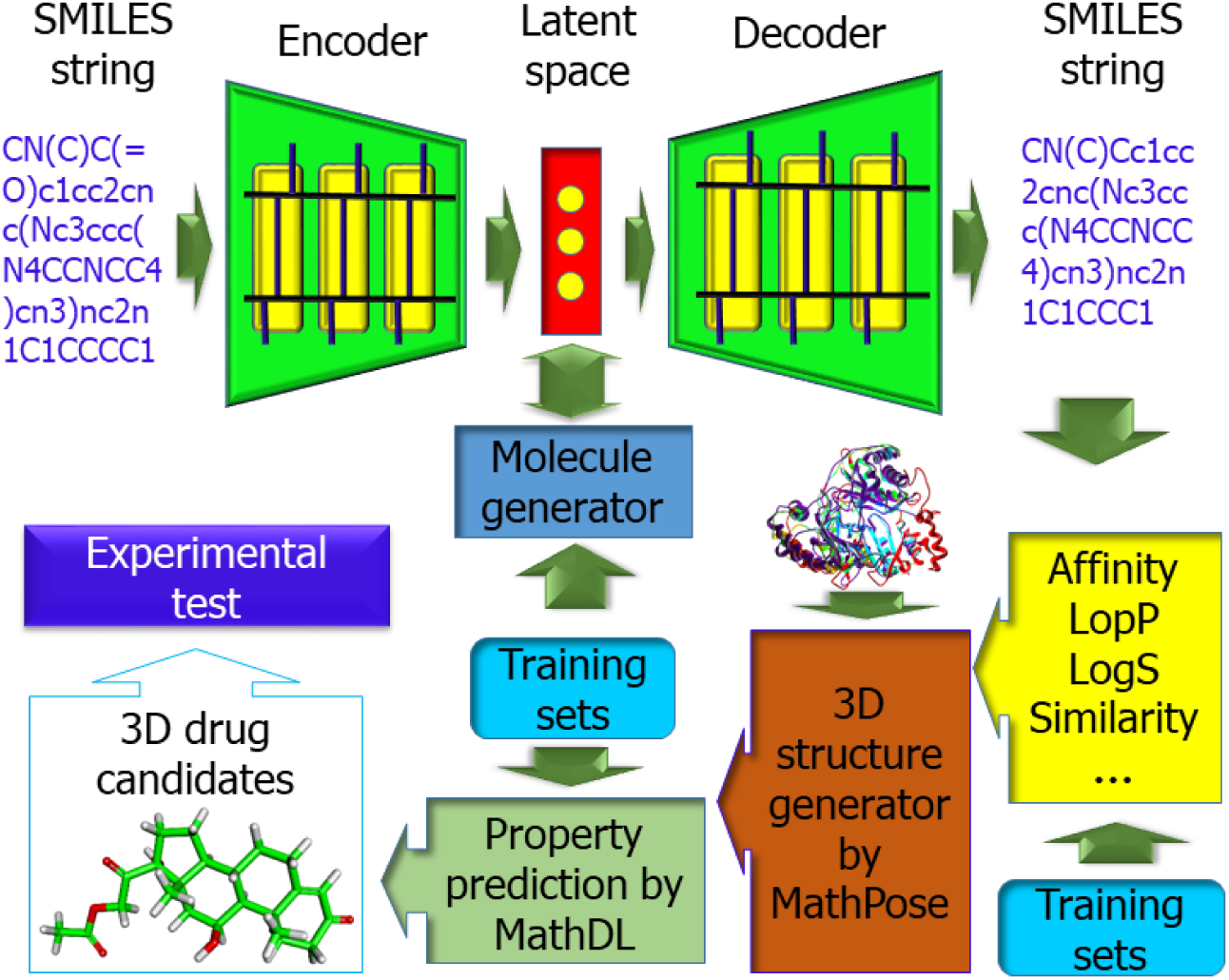
A schematic illustration of the generative network complex. SMILES strings (SSs) are encoded into latent space vectors via a gated recurrent neural network (GRU)-based encoder. These vectors are modified in the molecule generator to achieve desirable druggable properties, such as binding affinity, partition coefficient (LogP), similarity, etc., predicted by pre-trained deep neural networks (DNNs). The resulting drug-like molecules are translated into SSs by a GRU-based decoder. The physical properties of these SSs are validated by 2D fingerprint-based multitask DNNs. Promising drug candidates are fed into a MathPose unit to generate 3D structures, which are further validated by a mathematical deep learning (MathDL) center to select new drug candidates.^18^

### 2.1 Autoencoder

The autoencoder, consisting of an encoder, a latent space, and a decoder, is used to encode a molecular SMILES string into a latent space representation *X*, which, after being further modified by a molecular generator, is translated back to a SMILES string by a decoder. Both the encoder and decoder are constructed by using gated recurrent neural networks (GRUs). GRUs can deal with the vanishing gradient problem occurred in recurrent neural network (RNN) models but are simpler than long-short-term memory (LSTM) models. GRUs are suitable for moderately complex sequences, such as small molecular SMILES strings. A pre-trained autoencoder model developed by Winter et al is adopted in the present work.^19^ The latent space vector (*X* ∈ ℝ^*n*^) or molecular representation has the dimension of 512 (*n* = 512).

### 2.2 Molecule generator

In the present approach, novel molecules are initially designed at the molecule generator using a three-step procedure. In the first step, the latent space representation of a seed molecule is evaluated for their drug-like properties, such as binding affinities, solubility (logS), partition coefficient, similarity, etc., via pre-trained DNNs. The DNN consists of two hidden layers with 1024 neurons in each layer. In our second step, evaluation results ({ŷ_*i*_(*X*)}) are compared with a set of target values ({*y*_*i*0_}) via a loss function,

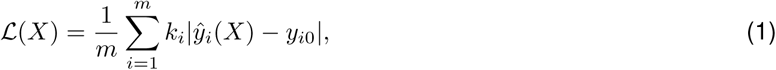

where *k*_*i*_ is a preselected weight coefficient for the *i*th property. The last step is to optimize the loss function with a gradient decent algorithm. The resulting new vector *X* is sent back to the pre-trained DNNs for reevaluation until the loss function is smaller than a given tolerance. Only the desirable molecule representation *X* is sent to the decoder to generate a SMILES string. Alternatively, a Monte Carlo procedure can be used to replace the gradient decent.

### 2.3 2D fingerprint-based predictor (2DFP)

New SMILES strings generated by the decoder is passed to 2D fingerprint-based predictors (2DFPs) to reevaluate druggable properties.^17^ These predictors are pre-trained deep neural networks involving multiple hidden layers with hundreds or even thousands of neurons on each layer. During the training, weights on each layer are updated by backpropagation. The multitask deep learning architecture is often used to enhance small dataset predictions. The input 2D molecular fingerprints are generated from a combination of ECFP^20^ and MACCS^21^ fingerprints, yielding 2214 bits of features (2048 bits from ECFP and 166 bits from MACCS) in total. RDKit^22^ is used for to translate SMILES strings into 2D fingerprints. The output drug properties include binding affinity, logP, and logS, etc.

### 2.4 MathDL for druggable property predictions

Our MathDL is a mathematical representation-based deep learning platform designed for predicting various druggable properties of 3D molecules.^18^ Mathematical representations used in MathDL are algebraic topology (such as persistent homology), differential geometry, and graph theory-based algorithms developed over the past many years. These approaches were repeatedly validated by their top performance in free energy prediction and ranking at D3R Grand Challenges, a worldwide competition series in computer-aided drug design (https://drugdesigndata.org/about/grand-challenge).^18,23^ More details about the mathematical representation of complex molecules can be found in a recent review.^24^ A variety of datasets, particularly, PDBbind datasets,^25^ were used in our training of deep learning networks. A further discussion of MathDL is given in our recent work.^17^

### 2.5 MathPose for 3D structure prediction

MathPose is a 3D pose predictor that converts SMILES strings into 3D poses with references of target molecules. For a given SMILES string, about 1000 3D structures are generated by several common docking software tools, i.e., Autodock Vina,^26^ GOLD,^27^ and GLIDE.^28^ Additionally, a selected set of known complexes is re-docked by the aforementioned three docking software packages to generate at 100 decoy complexes per input ligand as a machine learning training set. In this training set, the calculated root mean squared deviations (RMSDs) between the decoy and native structures are used as machine learning labels. Then, we set up MathDL models and apply them to pick up the top-ranked pose for the given ligand. The MathPose-generated top poses are fed to the MathDL for druggable property evaluation. Our MathPose was the top performer in D3R Grand Challenge 4 in predicting the poses of 24 beta-secretase 1 (BACE) binders.^18^

## 3 Results

### 3.1 Sequence identity analysis

The sequence identity is defined as the percentage of characters which match exactly between two different sequences. The sequence identities between 2019-nCoV protease and some other coronaviral proteases are presented in Table 1. It is seen that 2019-nCoV protease is very close to SARS-CoV protease, but is distinguished from other proteases. Clearly, 2019-nCoV has a strong genetic relationship with SARS-CoV. Additionally, the available experimental data of SARS-CoV protease inhibitors can be used as the training set to generate new inhibitors of 2019-nCoV protease.

**Table 1:**
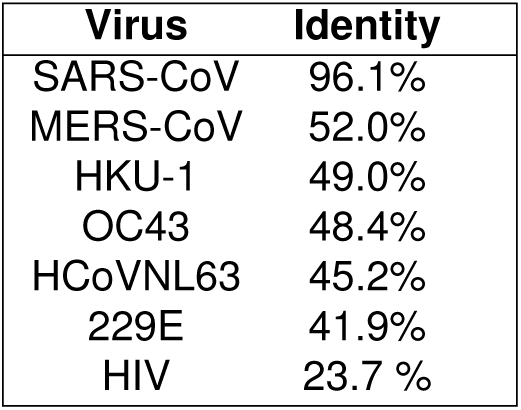
The sequence identity of the 2019-nCoV protease with some other viral proteases. The sequences of comparison are extracted from nCoV2019: https://www.ncbi.nlm.nih.gov/nuccore/MN908947, SARS: http://www.rcsb.org/structure/3IWM, MERS: http://www.rcsb.org/structure/5C3N, OC43: https://www.ncbi.nlm.nih.gov/protein/AEN19363.1, HCoVNL63: https://www.ncbi.nlm.nih.gov/nuccore/NC_005831, HKU-1: https://www.ncbi.nlm.nih.gov/nuccore/85667876, 229E: http://www.rcsb.org/structure/2ZU2, and HIV: http://www.rcsb.org/structure/2ZU2, and HIV: http://www.rcsb.org/structure/1YT9. Alignment is carried out by using https://web.expasy.org/sim/.

| Virus | Identity |
| --- | --- |
| SARS-CoV | 96.1% |
| MERS-CoV | 52.0% |
| HKU-1 | 49.0% |
| OC43 | 48.4% |
| HCoVNL63 | 45.2% |
| 229E | 41.9% |
| HIV | 23.7 % |

### 3.2 Structure similarity analysis

The 2019-nCoV protease (PDB ID 6lu7) and SARS-CoV 3CL protease (PDB ID: 2gx4) have an excellent similarity. As shown in Fig. 2, two crystal structures are essentially identical to each other. Particularly, the RMSD of two crystal structures at the binding site is 0.53 Å. When we try to carry a homology modeling of 2019-nCoV protease structure from its sequence using SARS-CoV 3CL protease (PDB ID: 2gx4) as a model, the resulting 2019-nCoV protease homology structure has an RMSD of 0.9 Å (or 0.2 Å at the binding site region) with its crystal structure 6lu7. The high structural similarity between the two proteases suggests that anti-SARS-CoV chemicals can be equally effective for the treatment of 2019-nCoV.

**Figure 2:**
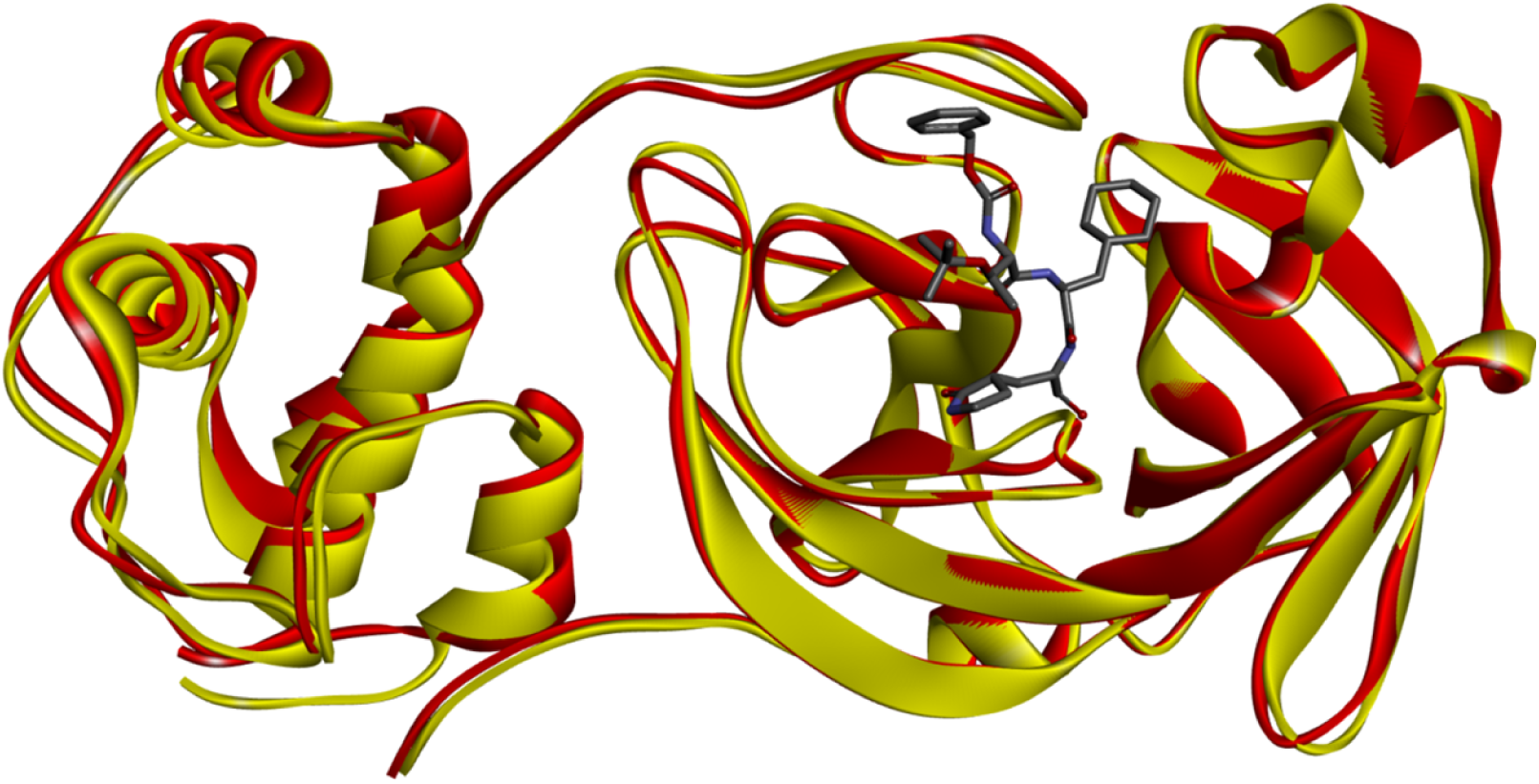
Illustration of the similarity between 2019-nCoV protease (PDB ID 6lu7) in gold and SARS-CoV 3CL protease (PDB ID: 2gx4) in red. The anti-SARS inhibitor in dark color indicates the binding site.

### 3.3 Datasets

#### 3.3.1 SARS-CoV protease inhibitor dataset

ChEMBL,^29^ an open database which brings chemical, bioactivity, and genomic data together to translate genomic information into effective new drugs is employed to construct our 2019-nCoV training set. Considering the high sequence identity between viral proteases of 2019-nCoV and SARS-CoV, we take the protease of SARS-CoV as the input target in ChEMBL and a total 115 ChEMBL IDs of the target can be found. Therefore, our 2019-nCoV training set is built up by 115 SARS-CoV protease inhibitors. Figure 4 describes the distribution of experimental Δ*G* values of 2019-nCoV training set. It can be seen that the experimental Δ*G* ranges from −10.0 kcal/mol to 7.5 kcal/mol, and most training samples have the experimental Δ*G* located in the range of [−10, −5] kcal/mol. Followed by the second law of thermodynamics, the more negative Δ*G* can result in a more spontaneous binding process. Figure 3 depicts the top five anti-SARS CoV compounds and their banding affinities.^29^ In this work, 115 SARS-nCoV protease inhibitors from ChEMBL are used as the 2019-nCoV training set. A collection of these 115 compounds is given in the Supplementary material.

**Figure 3:**
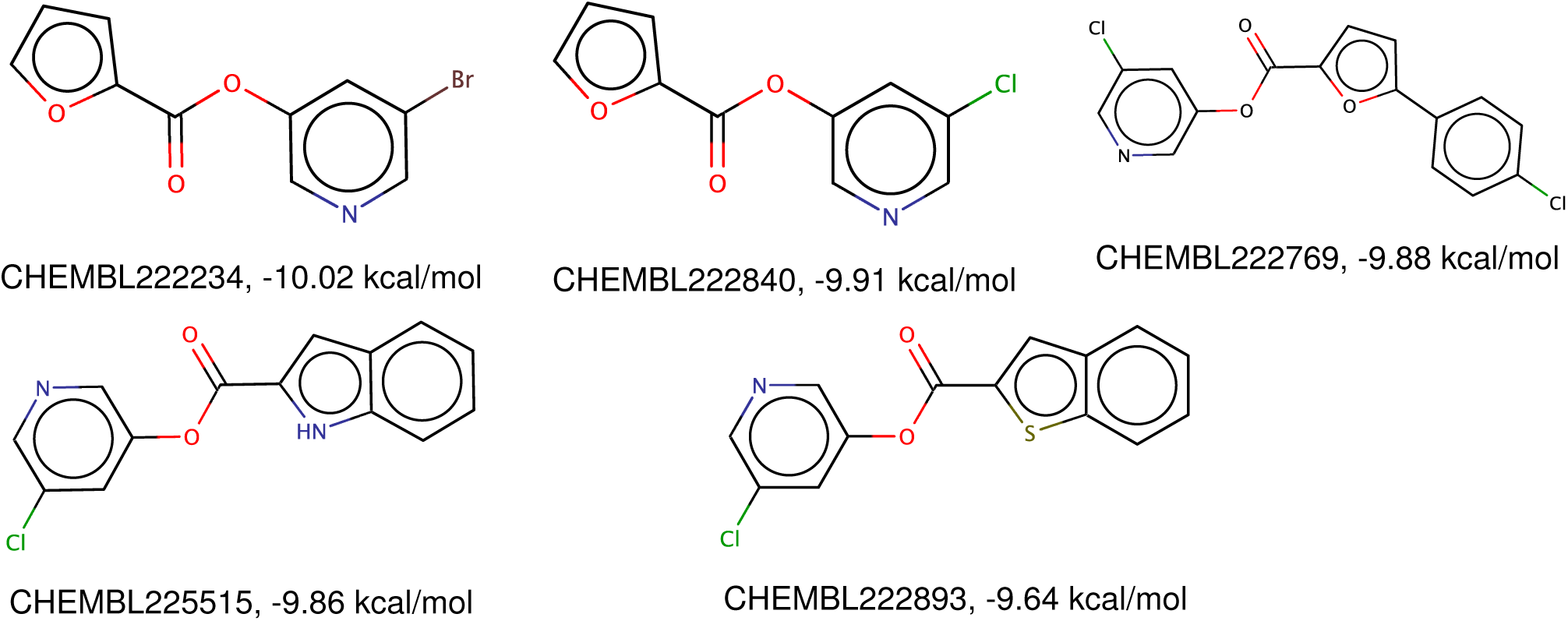
Top five anti-SARS-CoV compounds extracted from the ChEMBL database.

**Figure 4:**
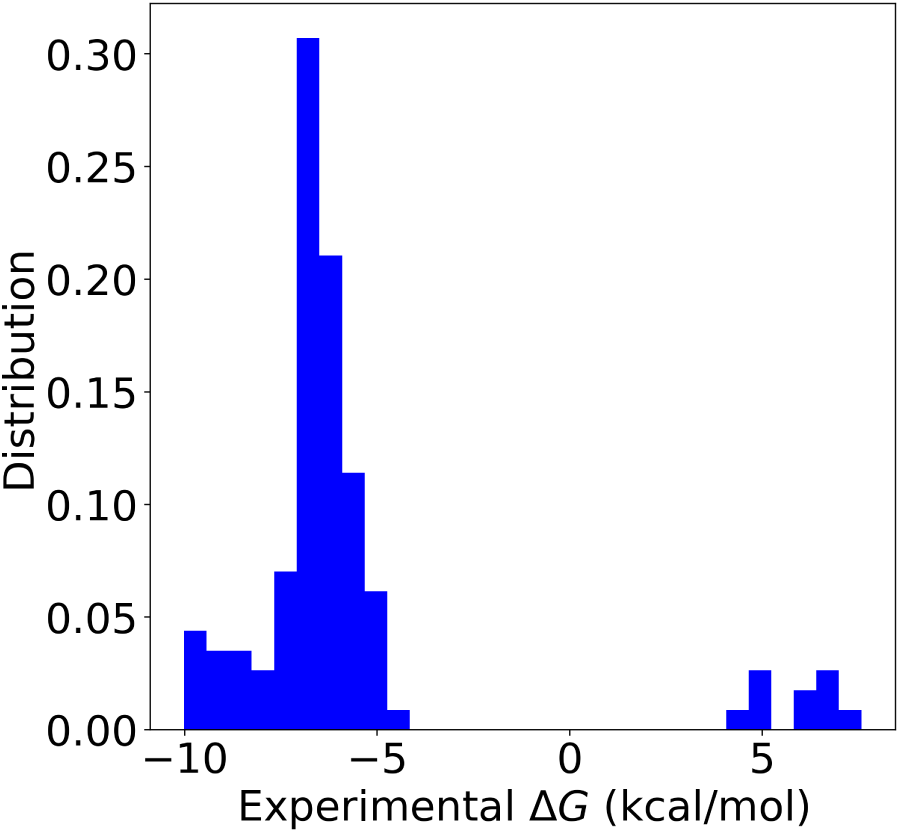
The distribution of experimental binding affinity values of SARS-CoV protease inhibitors.

#### 3.3.2 Binding affinity training sets

The PDBbind database is a yearly updated collection of experimentally measured binding affinity data (Kd, Ki, and IC50) for the protein-ligand complexes deposited in the Protein Data Bank (PDB). PDBbind refined set contains high-quality X-ray crystal structures of protein-ligand complexes and associated binding affinities.^25^ The refined set is selected by filtering through binding data, crystal structures, as well as the nature of the complexes.^25^ The PDBbind 2018 refined set of 4463 complexes is employed as the major part of our binding affinity training set. Additionally, taring data are collected from recent D3R Grand Challenges (https://drugdesigndata.org/about/grand-challenge) and a few hundreds of Pfizer molecules, which, however, mostly unrelated to the present coronaviral protease.

### 3.4 Binding affinity analysis

Binding free energies are computed with four methods, namely, the latent space binding predictor (LS-BP), the 2D fingerprint predictor (2DFP), a 3D deep learning model trained with the mixture all datasets, including the dataset for coronaviral protease (denoted as “3DALL”), and finally, a 3D deep learning multitask model trained with the dataset for coronaviral protease as a separated task (denoted as 3DMT).

Figures 5, 6, 7, 8, 9 10, 11, 12, 13, 14 15, 16, 17, 18 and 19 display GNC generated top 15 molecules. Their predicted binding affinities are given, together with their complexes with 2019-nCoV protease. These compounds are ranked according to their binding affinity values predicted by 3DALL scores. Predictions of other methods are also reported in Table 2 as references. Table 2 lists a few other druggable properties, including partition coefficient (logP), solubility (logS), and synthesizability.

**Table 2:**
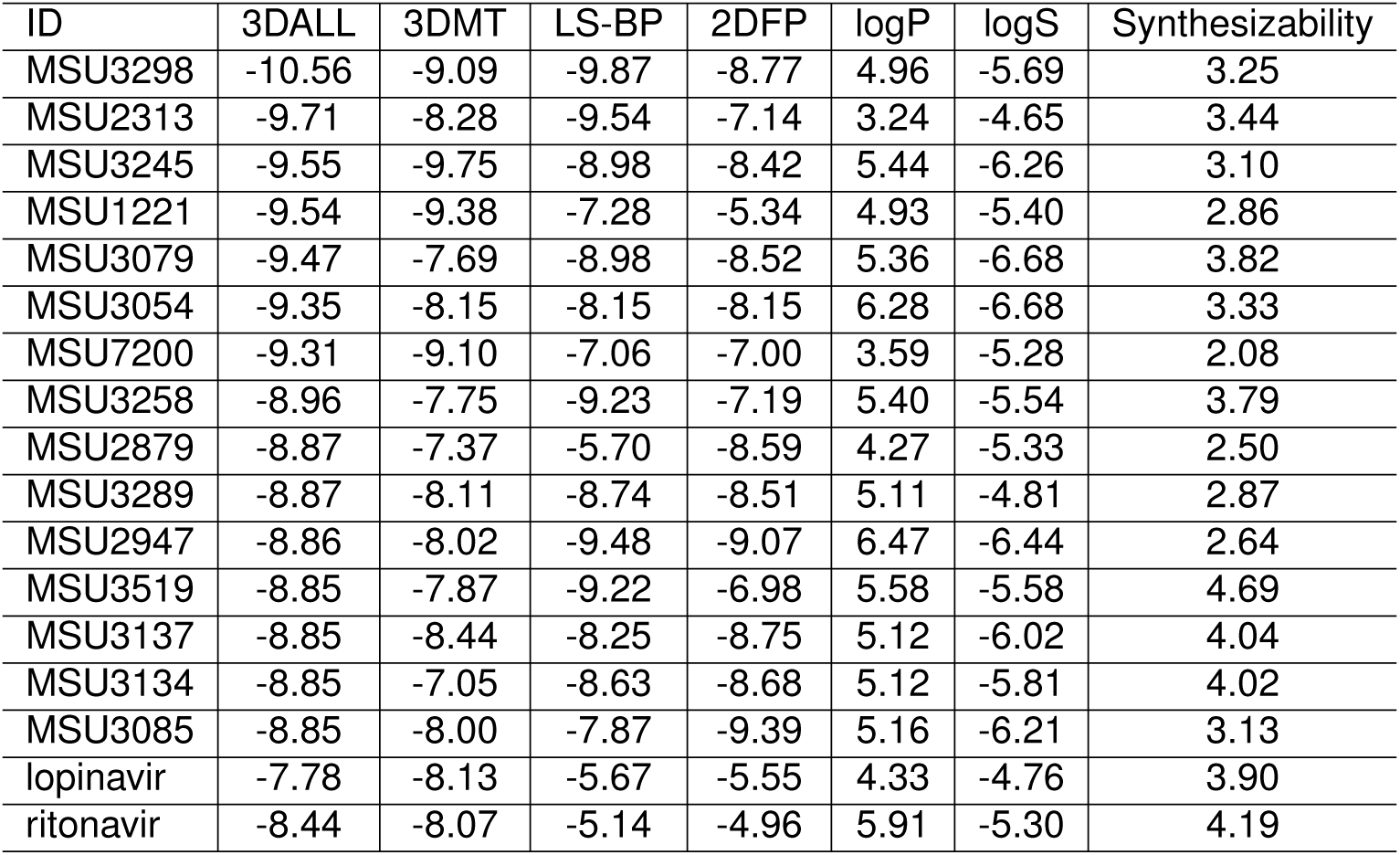
A summary of some druggable properties for the top 15 anti-2019-nCoV molecules generated by GNC and two HIV drugs.

| ID | 3DALL | 3DMT | LS-BP | 2DFP | logP | logS | Synthesizability |
| --- | --- | --- | --- | --- | --- | --- | --- |
| MSU3298 | -10.56 | -9.09 | -9.87 | -8.77 | 4.96 | -5.69 | 3.25 |
| MSU2313 | -9.71 | -8.28 | -9.54 | -7.14 | 3.24 | -4.65 | 3.44 |
| MSU3245 | -9.55 | -9.75 | -8.98 | -8.42 | 5.44 | -6.26 | 3.10 |
| MSU1221 | -9.54 | -9.38 | -7.28 | -5.34 | 4.93 | -5.40 | 2.86 |
| MSU3079 | -9.47 | -7.69 | -8.98 | -8.52 | 5.36 | -6.68 | 3.82 |
| MSU3054 | -9.35 | -8.15 | -8.15 | -8.15 | 6.28 | -6.68 | 3.33 |
| MSU7200 | -9.31 | -9.10 | -7.06 | -7.00 | 3.59 | -5.28 | 2.08 |
| MSU3258 | -8.96 | -7.75 | -9.23 | -7.19 | 5.40 | -5.54 | 3.79 |
| MSU2879 | -8.87 | -7.37 | -5.70 | -8.59 | 4.27 | -5.33 | 2.50 |
| MSU3289 | -8.87 | -8.11 | -8.74 | -8.51 | 5.11 | -4.81 | 2.87 |
| MSU2947 | -8.86 | -8.02 | -9.48 | -9.07 | 6.47 | -6.44 | 2.64 |
| MSU3519 | -8.85 | -7.87 | -9.22 | -6.98 | 5.58 | -5.58 | 4.69 |
| MSU3137 | -8.85 | -8.44 | -8.25 | -8.75 | 5.12 | -6.02 | 4.04 |
| MSU3134 | -8.85 | -7.05 | -8.63 | -8.68 | 5.12 | -5.81 | 4.02 |
| MSU3085 | -8.85 | -8.00 | -7.87 | -9.39 | 5.16 | -6.21 | 3.13 |
| lopinavir | -7.78 | -8.13 | -5.67 | -5.55 | 4.33 | -4.76 | 3.90 |
| ritonavir | -8.44 | -8.07 | -5.14 | -4.96 | 5.91 | -5.30 | 4.19 |

**Figure 5:**
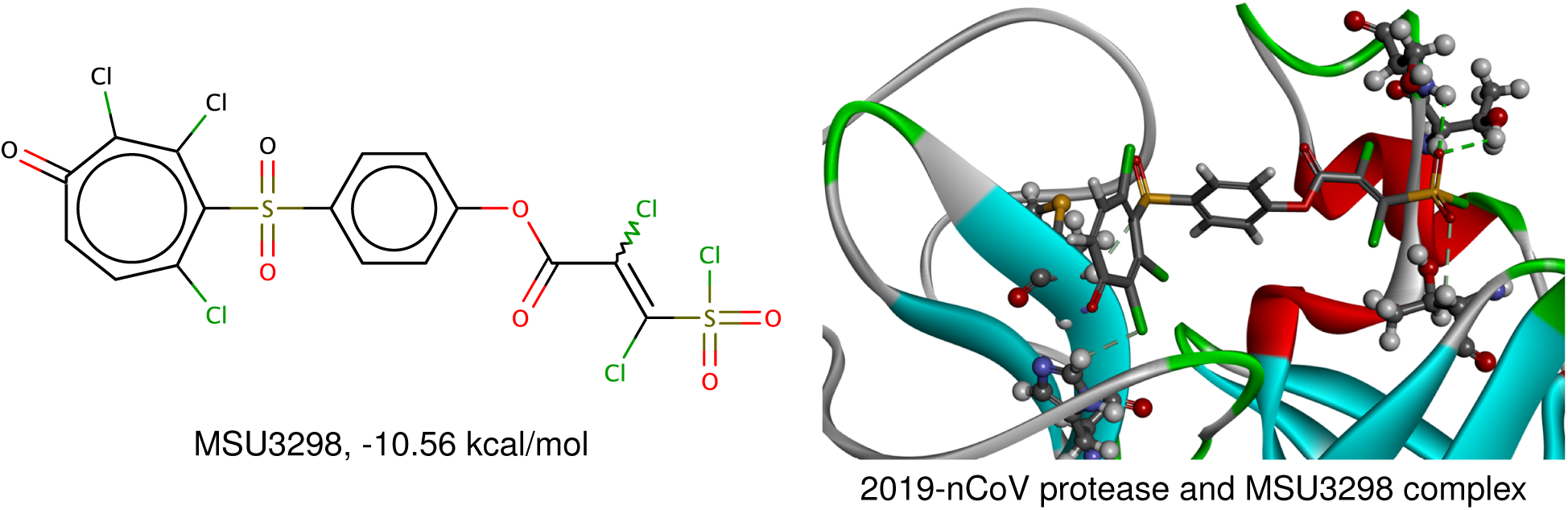
MSU3298 molecule and its complex with 2019-nCoV protease.

**Figure 6:**
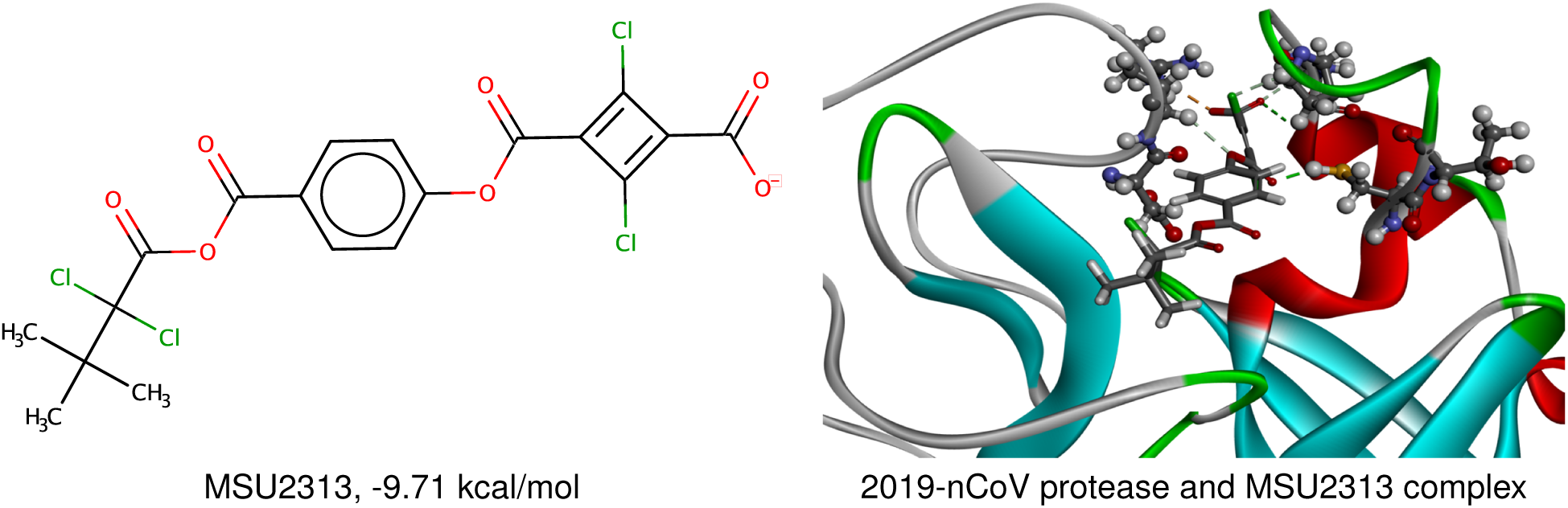
MSU2313 molecule and its complex with 2019-nCoV protease.

**Figure 7:**
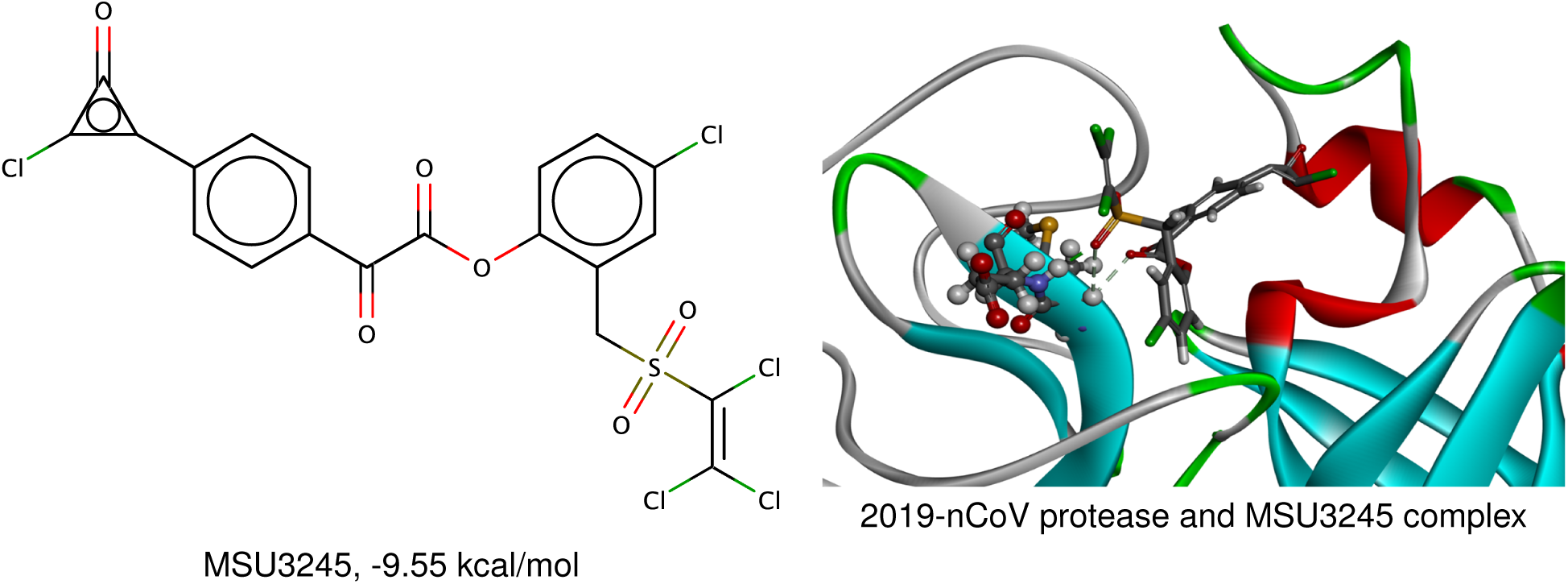
MSU3245 molecule and its complex with 2019-nCoV protease.

**Figure 8:**
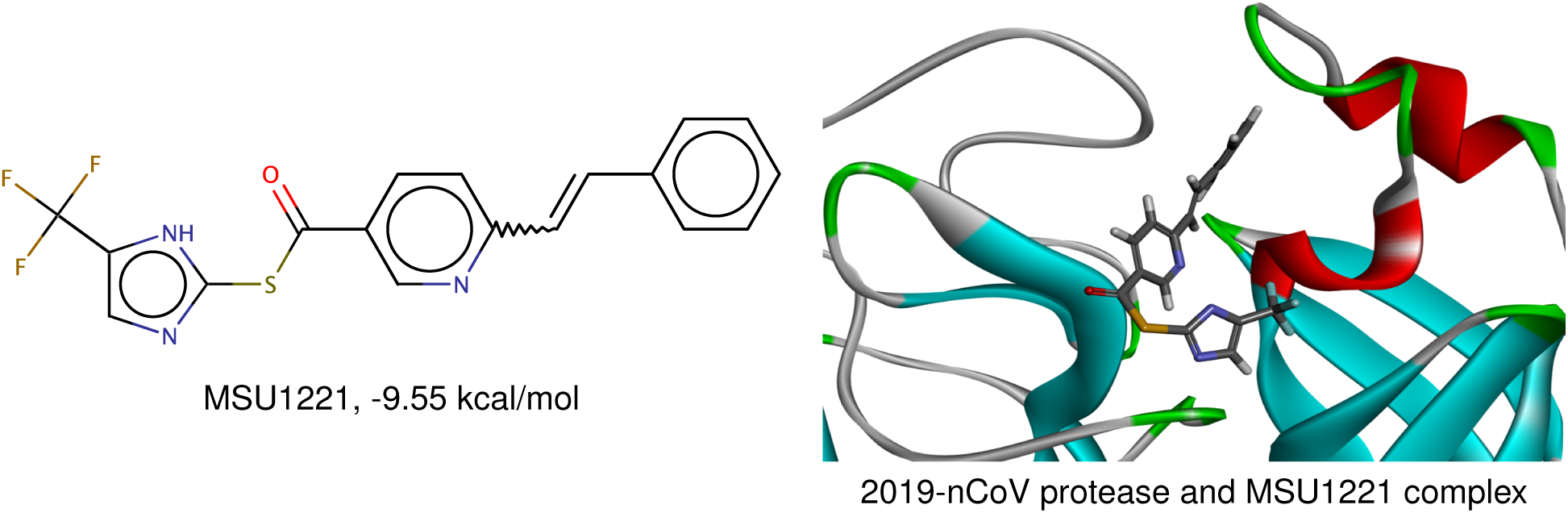
MSU1221 molecule and its complex with 2019-nCoV protease.

**Figure 9:**
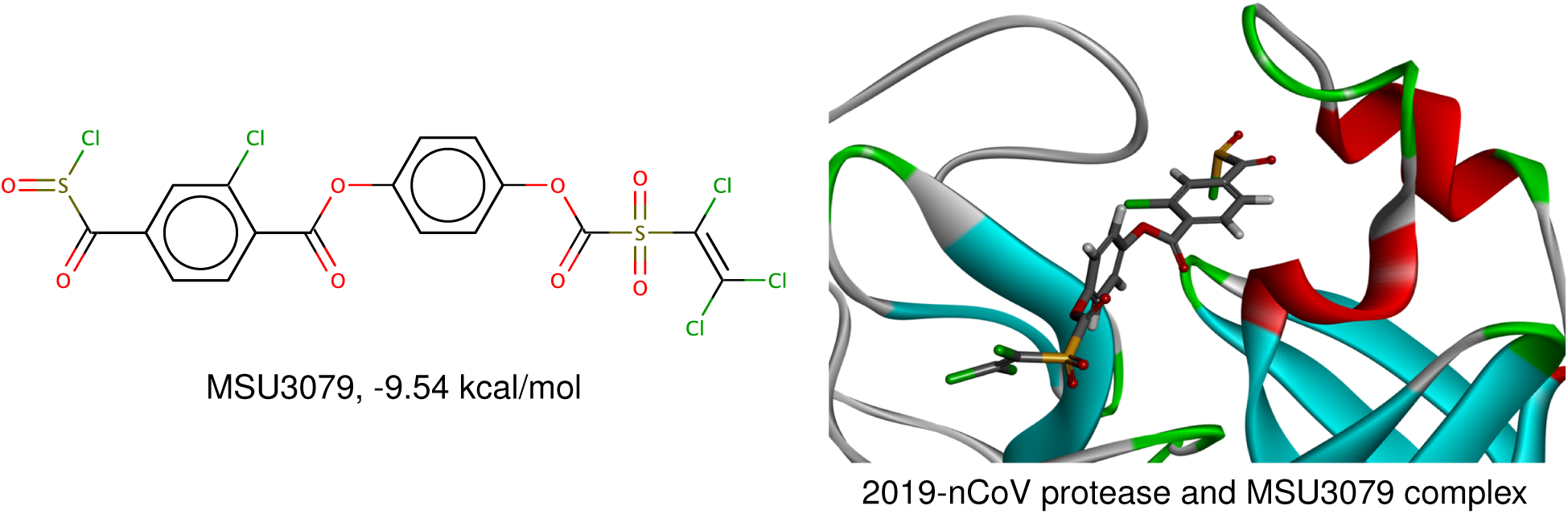
MSU3079 molecule and its complex with 2019-nCoV protease.

**Figure 10:**
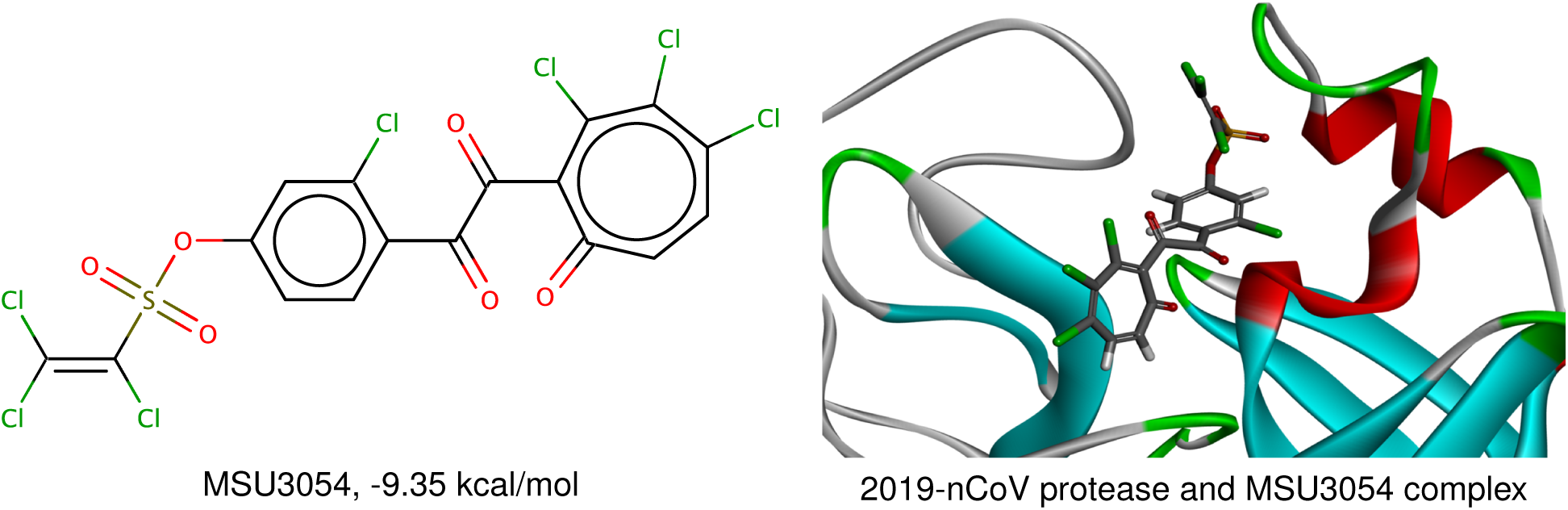
MSU3054 molecule and its complex with 2019-nCoV protease.

**Figure 11:**
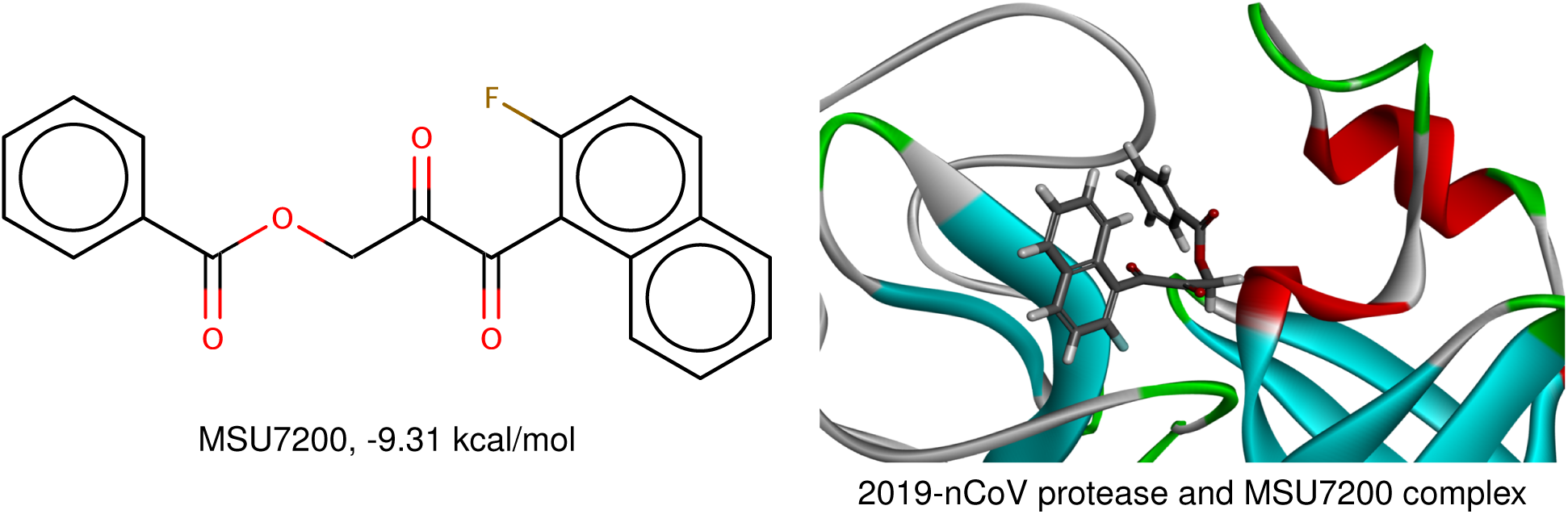
MSU7200 molecule and its complex with 2019-nCoV protease.

**Figure 12:**
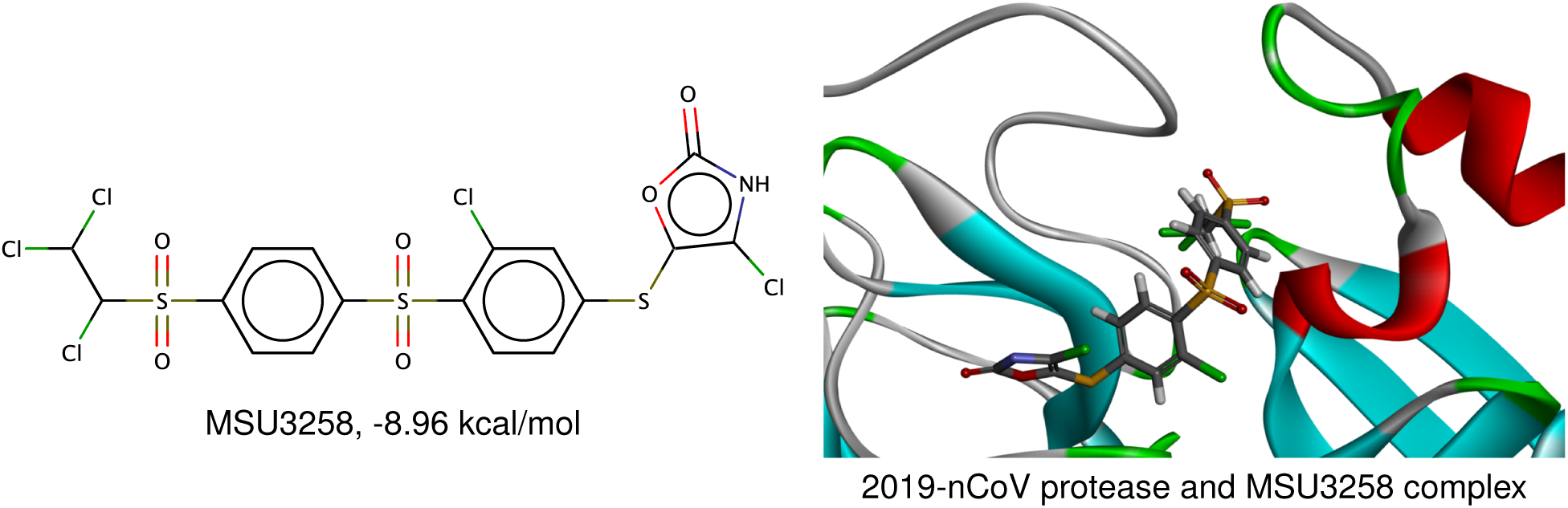
MSU3258 molecule and its complex with 2019-nCoV protease.

**Figure 13:**
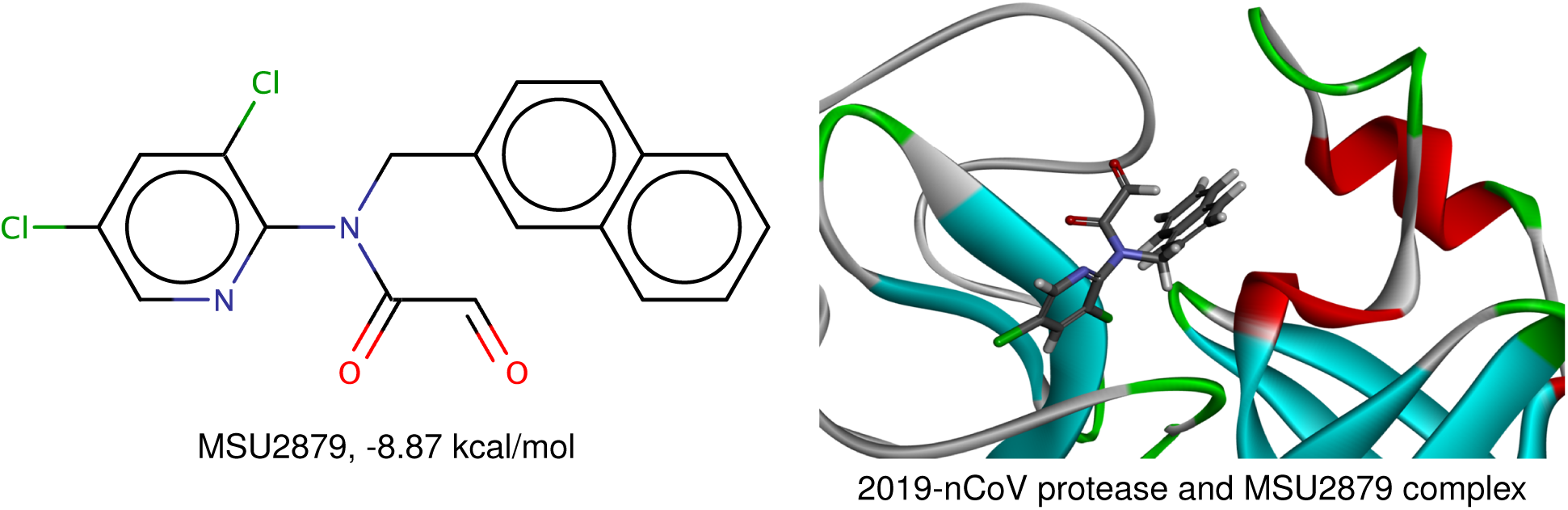
MSU2879 molecule and its complex with 2019-nCoV protease.

**Figure 14:**
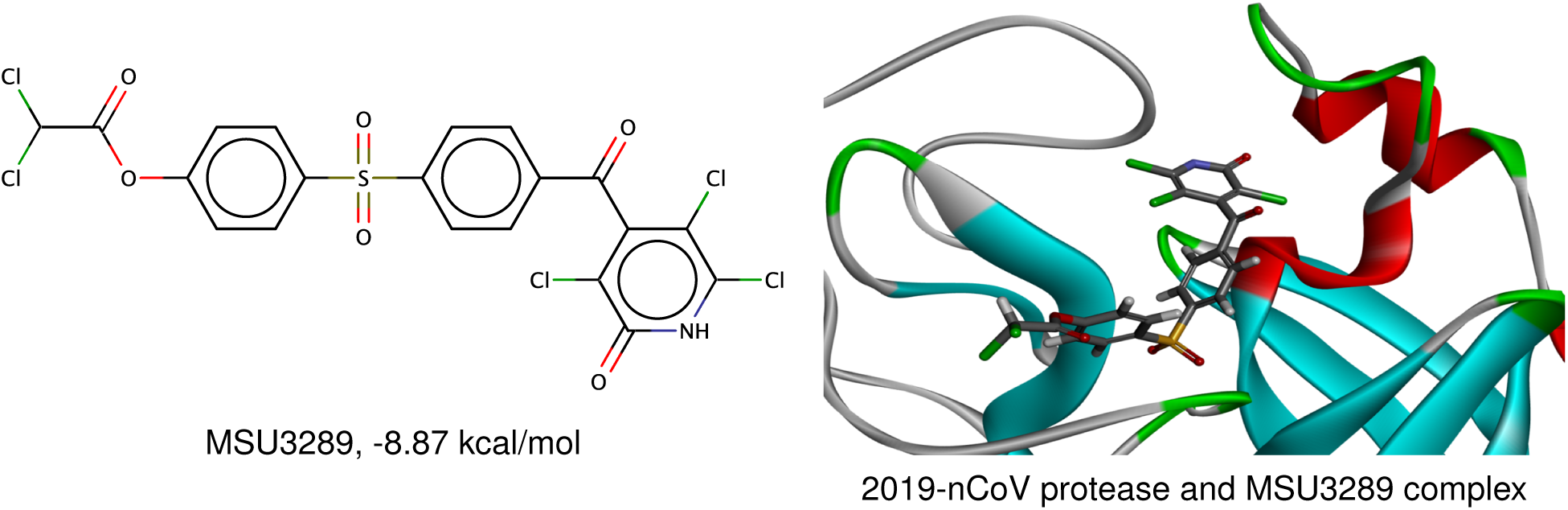
MSU3289 molecule and its complex with 2019-nCoV protease.

**Figure 15:**
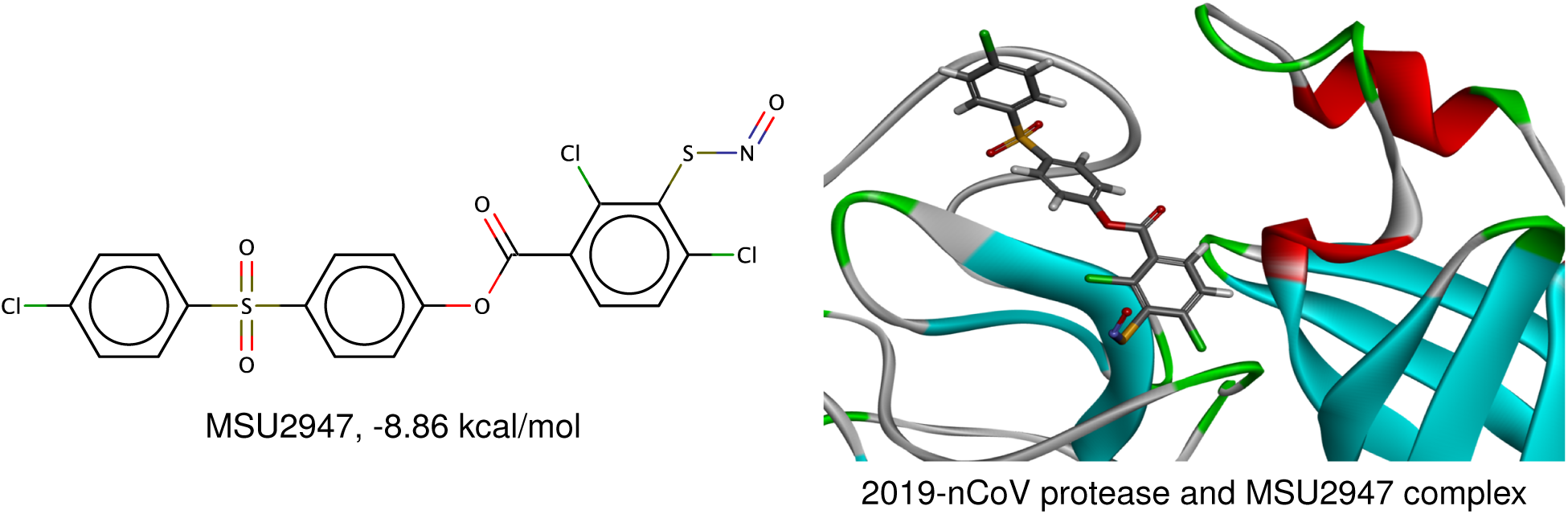
MSU2947 molecule and its complex with 2019-nCoV protease.

**Figure 16:**
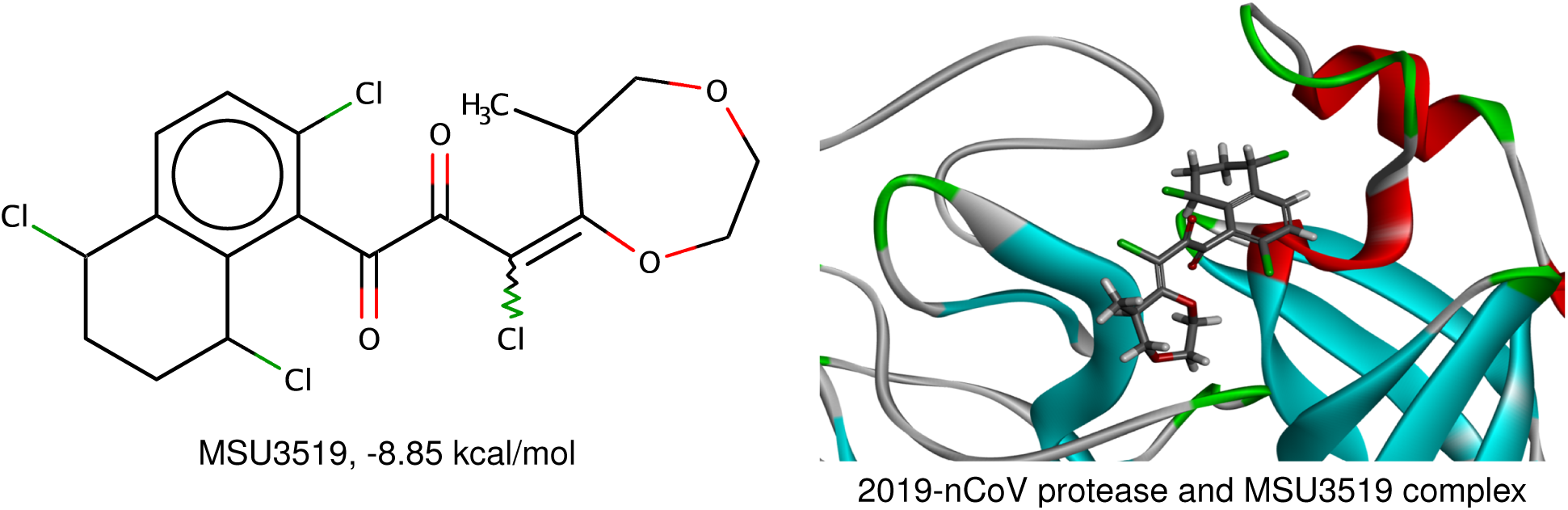
MSU3519 molecule and its complex with 2019-nCoV protease.

**Figure 17:**
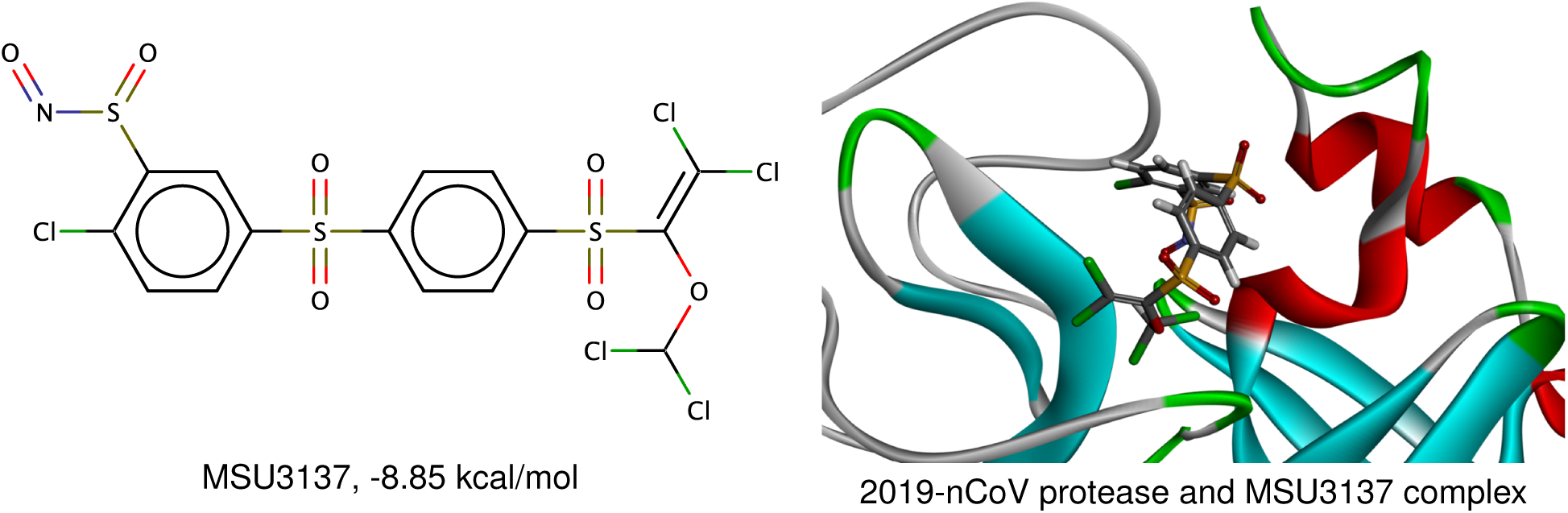
MSU3137 molecule and its complex with 2019-nCoV protease.

**Figure 18:**
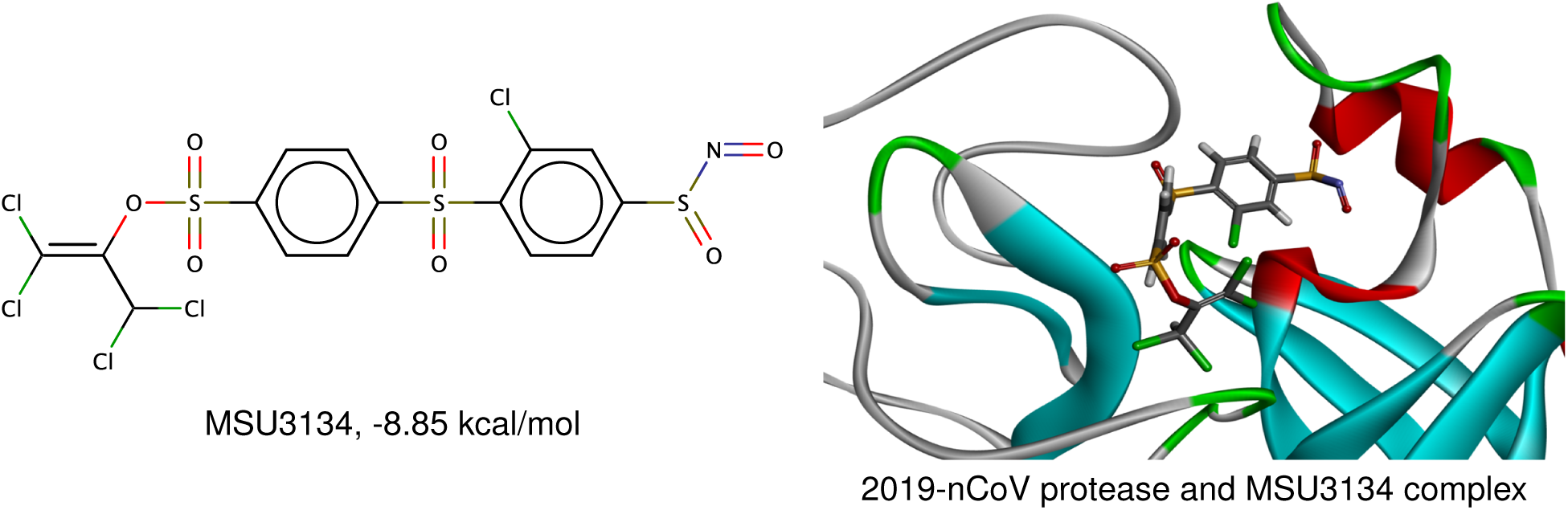
MSU3134 molecule and its complex with 2019-nCoV protease.

**Figure 19:**
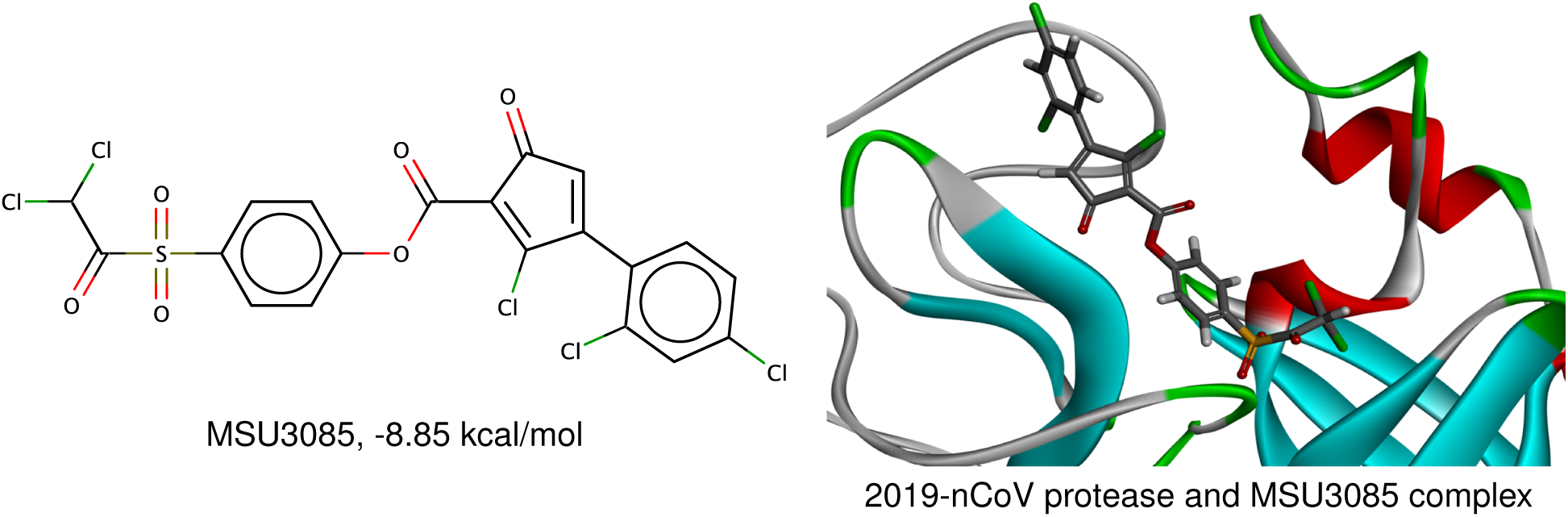
MSU3085 molecule and its complex with 2019-nCoV protease.

The top-ranking candidate of our generated molecules is MSU3298 (see Figure 5). Its predicted binding affinity to the nCoV-2019 protease is −10.56 kcal/mol, which is higher than that of the best candidate CHEMBL222234 (−10.02 kcal/mol). The high binding affinity is due to the existence of many hydrogen bond acceptors and donors and forming a strong hydrogen bond network with nCoV-2019 protease. For example, the strongest hydrogen MSU3245, −9.55 kcal/mol bonds are formed by the three Cl atoms on the tail of MSU3298 molecule and three different Hydroxyls in the residues Thr45, Ser46 and Thr25 of nCoV-2019 protease. This tail bonds tightly with the side chains of the aforementioned residues. Another important interaction is located at the head of the molecule. A hydrogen bond formed between the Cl atom in the heptatomic ring and the Hydroxyl in the sidechain of residue Ser144. Moreover, one O atom in the methylsulfonylmethane of the molecule also forms a hydrogen bond with the residue Met165. As a result, the head, body, and tail of the molecule interact firmly with the protease inhibition binding site.

For the second-best molecule, MSU2313 (see Figure 6), the binding affinity is −9.71 kcal/mol. The high binding affinity is also due to the strong interaction at the tail of the molecule. One Cl atom on the quaternary ring forms a hydrogen bond with the side chain of residue Arg188. One O atom in this tail also has a hydrogen bond with the main chain of residue Tyr54. In the head of the molecules, the two Cl atoms interact with the methanethiol of residue Cys145 and the main-chain amino of residue Glu166. These hydrogen bonds promise a strong binding to 2019-nCoV protease binding site.

The third molecule is MSU3245 (see Figure 7) with a binding affinity −9.55 kcal/mol. The strongest hydrogen bonds between this molecule and the protease are the Cl atom on the benzene ring of the molecule and the side-chain hydroxyl of the residue Ser144. Additionally, the Cl atom of the ternary ring and the methanethiol in residue Cys144.

Essentially, the hydrogen bond acceptor such as Cl and O atoms in the drug candidate molecules are critical to the binding to 2019-nCoV protease. The hydrogen bond acceptors can form strong hydrogen bonds with the nCoV-2019 protease and inhibit its function.

## 4 Discussions

### 4.1 Solubility

Aqueous solubility, a chemical property denoted by its logarithm value logS, reveals how a solute dissolves in a solvent which will affect absorption, distribution, metabolism, and elimination processes (ADME) in drug discovery and other pharmaceutical fields.^30^ It is part of the pharmacokinetics studies. In this work, we calculate the logS values for all of the new potential anti-2019-nCoV drugs using our 2DFP-based logS predictor. Table 2 lists top 15 anti-2019-nCoV candidate molecules with their druggable properties. It is seen that the smallest logS is −6.44 and the largest value is −4.65. According to the literature,^31,32^ about 85% of drugs have logS values between −5.000 and −1.000. However, only two potential anti-2019-nCoV drugs (i.e., MSU2313 and MSU3289) in Table 2 have the logS values in the range of [−5.00, −1.00], while the others have a little bit higher value of logS. One possible reason is that our 2DFP-based calculation of logS may have a systematic error. Another possible explanation is that our anti-2019-nCoV drug candidates may not be absorbed through membranes as easily as some other drugs on the market. In our future study, a stronger logS constraint will be imposed in our molecule generator.

### 4.2 Partition coefficient

The partition coefficient, which measures how hydrophilic or hydrophobic a chemical substance is, is defined as the ratio of concentrations of a solute in a mixture of two immersible solvents at equilibrium.^33^ The logarithm of partition coefficient, denoted logP, is a well-known coefficient which plays an essential role in governing kinetic and dynamic aspects of drug action. In this paper, we will employ an Open-Source Cheminformatics Software Rdkit^22^ to calculate logP values of our 2019-nCoV drug candidates to evaluate the reliability of the potential 2019-nCoV drugs we predicted. All of the logP values of all predicted molecules can be found in the Supplementary Materials. While the logP values of the predicted top 15 drug candidates are presented in Table 2. From the table, it can be observed that most 2019-nCoV drug candidates we predicted have the logP value smaller than 5, which matches one of the rules in “Lipinski’s rule of five”.^34^ Moreover, the ritonavir, an HIV protease inhibitor already on the market, has a predicted logP = 5.91, which shows that our potential drugs with logP values slightly larger than 5 can still be considered as druggable molecules.

Moreover, consider the fact that logP also has the ability to measure the solubility of the solute in liquids, we are more likely to say that our logS calculation method in subsection 4.1 is not as precise as expected. These issues can be addressed by placing a stronger logS constraint in the latent space.

### 4.3 Synthesizability

Although we have the chemical structures of possible anti-2019-nCoV drugs, it is essential for us to estimate the feasibility of synthesis (synthetic accessibility) of these molecules. The synthetic accessibility score (SAscore) between 1 (easy to make) and 10 (very difficult to make) is described in.^35^ The SAscores of our drug candidates are calculated by the Rdkit as the evaluation of the molecules’ synthesizability. The last column in the Table 2 lists the SAscores of our Top 15 anti-2019-nCoV molecules. The molecule ID: MSU3519 has the highest SAscore equals to 4.69, which reveals that most of our potential anti-2019-nCoV molecules are quite easy to synthesize.

### 4.4 Effectiveness of some anti-HIV/AIDS drugs for 2019-nCoV

Lopinavir is an antiretroviral medication used to inhibit HIV/AIDS viral protease. It is often used as a fixed-dose combination with another protease inhibitor, ritonavir, sold under the name Kaletra or Aluvia. Ritonavir, sold under the trade name Norvir, is another antiretroviral medication. Its combination with Lopinavir is known as highly active antiretroviral therapy (HAART). Although there is no tractable clinical evidence, Kaletra or Aluvia has been proposed as a potential anticoronavirus drug for 2019-nCoV. The possibility of repurposing some HIV drugs for SARS-CoV treatment has also studied in the literature.^16^ It is important to evaluate their binding affinities, which are obtained with two ligand-based methods (i.e., LS-BP and 2DFP) and two 3D models (3DALL and 3DMT). To carry out 3D model predictions, we dock them to the 2019-nCoV protease inhibition site. The resulting complexes are optimized with molecular dynamics and then evaluated by 3DALL and 3DMT.

Table 1 shows the low sequence identity between HIV viral protease and 2019-nCoV protease, which might suggest the limited potential for repurposing Aluvia and Norvir for 2019-nCoV treatment. For Lopinavir, our LS-BP and 2DFP predicted the binding affinities of −5.66 kcal/mol and −5.54 kcal/mol, respectively. For Ritonavir, similar low binding affinities of −5.14 kcal/mol and −4.96 kcal/mol were predicted by our LS-BP and 2DFP, respectively. However, our 3D model 3DALL predicted better binding affinities, i.e., −7.78 kcal/mol and −8.44 kcal/mol for Lopinavir and Ritonavir, respectively. The other 3D model, 3DMT, also predicted moderately high binding affinities of −8.13 kcal/mol and −8.07 kcal/mol for Lopinavir and Ritonavir, respectively. Considering the fact that the small training set for LS-BP and 2DFP models is very small, the results predicted by 3D models are more reliable. Figures 20 and 21 indicate that these drugs have reasonable dock poses with 2019-nCoV protease. Therefore, HIV drugs Kaletra (or Aluvia) and Norvir might indeed have a moderate effect in the treatment of 2019-nCoV. However, Many new compounds generated by our GNC appear to have better druggable properties than these HIV inhibitors do.

**Figure 20:**
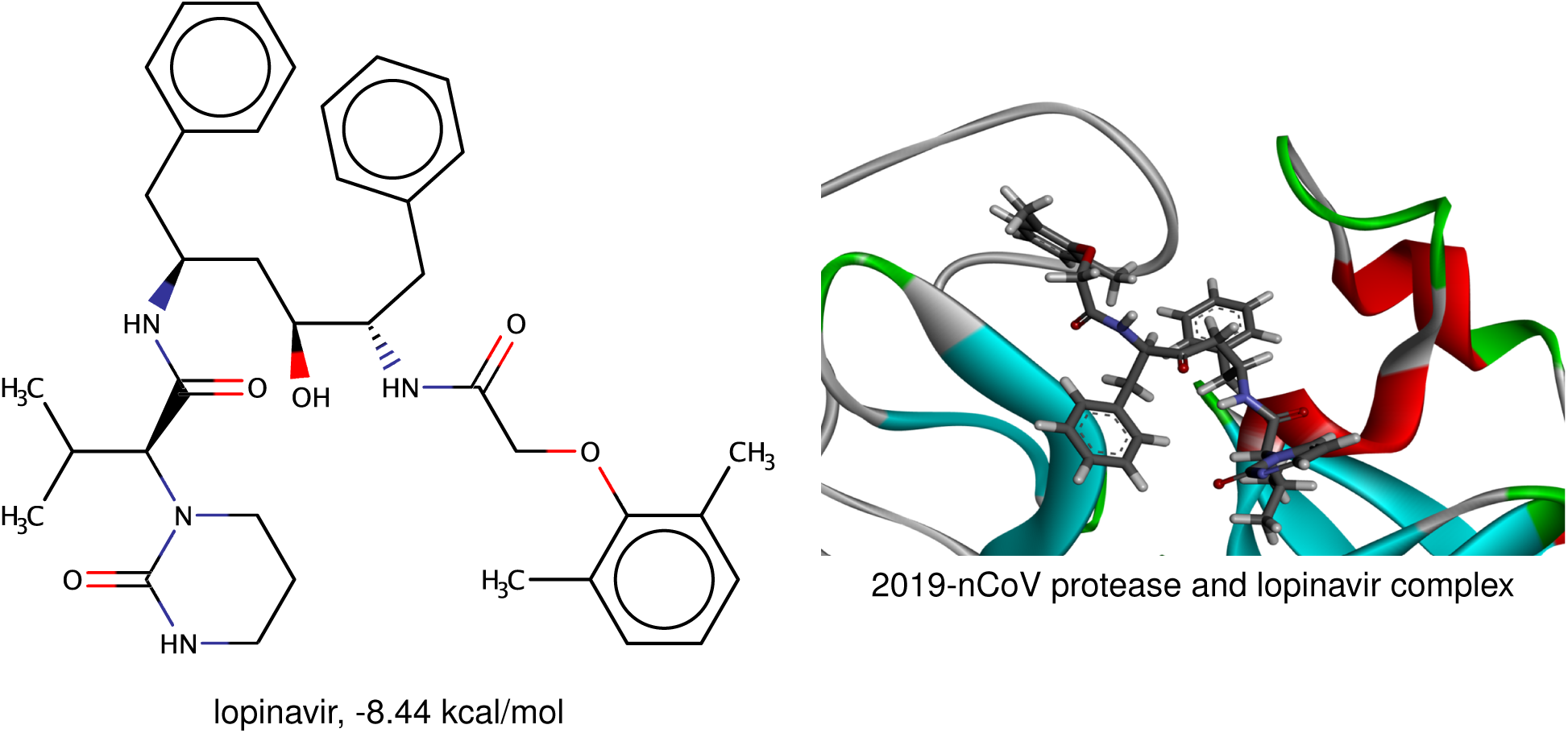
HIV drug lopinavir and its complex with 2019-nCoV protease. The complex structure showing a reasonable fit.

**Figure 21:**
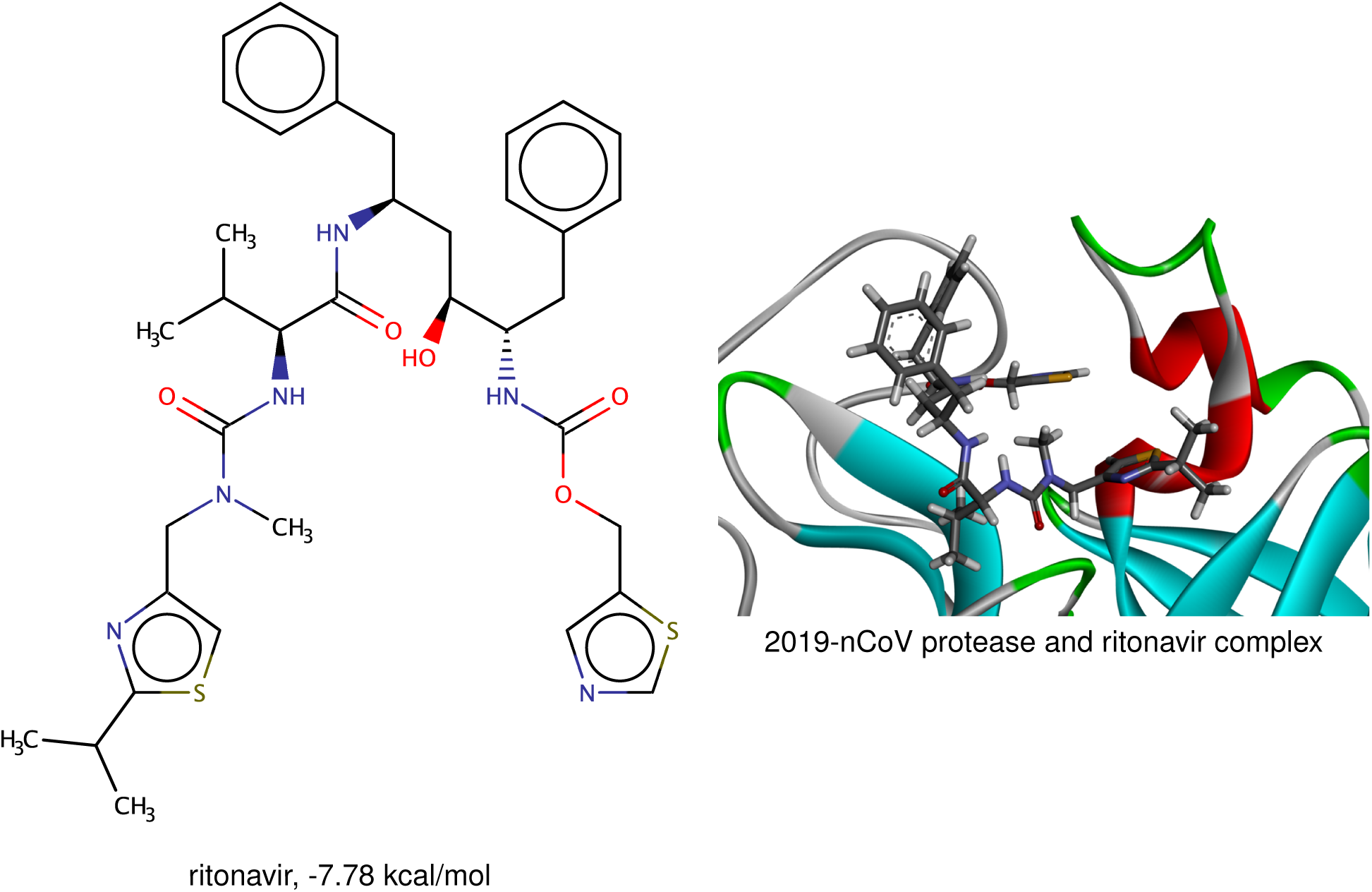
HIV drug ritonavir and its complex with 2019-nCoV protease. The complex structure showing a reasonable fit.

## 5 Conclusion

Wuhan pneumonia outbreak caused by a new coronavirus (CoV), called 2019-nCoV, has led to heavy economic loss and human fatalities. Under the current health emergency, it is vital to develop an effective treatment for this epidemic. Although we know quite a little about 2019-nCoV, it is fortunate that the sequence identity of the 2019-nCoV protease and that of severe acute respiratory syndrome virus (SARS-CoV) is as high as 96.1%. In this work, we show that the protease inhibitor binding sites of 2019-nCoV and SARS-CoV are almost identical, which provides a foundation for us to hypothesize that all potential anti-SARS-CoV chemotherapies are also effective anti-2019-CoV molecules. Additionally, we employed a recently developed generative network complex (GNC) to seek potential protease inhibitors for effective treatment of pneumonia caused by 2019-nCoV. Two datasets are utilized in this work. One is a SARS-CoV protease inhibitor dataset, which is constructed by collecting 115 SRAS-CoV inhibitors from open database ChEMBL. The other dataset is a binding affinity training set mainly containing the PDBbind refined set. Our GNC model predicts over 8000 potential anti-2019-nCoV drugs which are evaluated by a latent space binding predictor (LS-BP) and a 2D fingerprint predictor (2DFP). Promising drug candidates are further evaluated by two 3D deep learning models trained with all the training sets together, including the dataset for coronaviral protease (3DALL), and the 3D deep learning multitask model trained with the dataset for coronaviral protease as a separated task (3DMT). Furthermore, we choose 15 potential anti-2019-nCoV drugs to analyze partition coefficient (logP), solubility (logS), and synthetic accessibility score (SAscore) according to binding affinity ranking computed by the 3DALL model. The reasonable logP, logS, and SAscore show that our top 15 anti-2019-nCoV drug candidates are potentially effective for inhibiting 2019-nCoV. Finally, the effectiveness of some anti-HIV/AIDS drugs for treating 2019-nCoV is analyzed. Although HIV drugs Kaletra (or Aluvia) and Norvir might indeed have a moderate effect in the treatment of 2019-nCoV, the analysis of these anti-HIV/AIDS drugs together with our top 15 anti-2019-nCoV molecules shows that the new compounds generated by our GNC appear to have better druggable properties than these HIV inhibitors do.

## Supplementary materials

SupplementaryMaterial-1.csv: A list of SMILES strings, predicted binding affinities, logP, logS and synthesizability of 319 potential inhibitors of 2019-nCoV 3CL protease, including two anti-HIV protease drugs.

SupplementaryMaterial-2.csv: The SMILES strings, experimental binding affinities of 115 potential SARS inhibitors from ChEMBL. They are used as a training set for our GNC.

## Acknowledgments

This work was supported in part by NIH grant GM126189, NSF Grants DMS-1721024, DMS-1761320, and IIS1900473, Bristol-Myers Squibb, and Pfizer. The authors thank Zihe Rao & Haitao Yang group at Shanghai Technological University for providing the crystal structure of 2019-nCoV 3CL hydrolase (PDB ID 6lu7) and thank Dr. Min Li for helpful discussions.

## References

[1] Biao Tang, Xia Wang, Qian Li, Nicola Luigi Bragazzi, Sanyi Tang, and Yanni Xiao. Estimation of the transmission risk of 2019-ncov and its implication for public health interventions. SSRN, 2020.

[2] David S Hui, TA Madani, F Ntoumi, R Kock, O Dar, G Ippolito, TD Mchugh, ZA Memish, C Drosten, A Zumla, et al. The continuing 2019-ncov epidemic threat of novel coronaviruses to global health-the latest 2019 novel coronavirus outbreak in wuhan, china. International journal of infectious diseases: IJID: official publication of the International Society for Infectious Diseases, 91:264, 2020.

[3] Vincent CC Cheng, Shuk-Ching Wong, Kelvin KW To, PL Ho, and Kwok-Yung Yuen. Preparedness and proactive infection control measures against the emerging wuhan coronavirus pneumonia in china. Journal of Hospital Infection, 2020.

[4] Michael C Letko and Vincent Munster. Functional assessment of cell entry and receptor usage for lineage b *β*-coronaviruses, including 2019-ncov. bioRxiv, 2020.

[5] Xintian Xu, Ping Chen, Jingfang Wang, Jiannan Feng, Hui Zhou, Xuan Li, Wu Zhong, and Pei Hao. Evolution of the novel coronavirus from the ongoing wuhan outbreak and modeling of its spike protein for risk of human transmission. SCIENCE CHINA Life Sciences, 2020.

[6] Ning Dong, Xuemei Yang, Lianwei Ye, Kaichao Chen, Edward Wai-Chi Chan, Mengsu Yang, and Sheng Chen. Genomic and protein structure modelling analysis depicts the origin and infectivity of 2019-ncov, a new coronavirus which caused a pneumonia outbreak in wuhan, china. bioRxiv, 2020.

[7] Lisa E Gralinski and Vineet D Menachery. Return of the coronavirus: 2019-ncov. Viruses, 12(2):135, 2020.

[8] Yongzhen Zhang. Initial genome release of novel coronavirus. http://virological.org/t/novel-2019-coronavirus-genore/319/, 2020.

[9] Fang Li. Structure, function, and evolution of coronavirus spike proteins. Annual review of virology, 3:237–261, 2016.

[10] Lanying Du, Richard Y Kao, Yusen Zhou, Yuxian He, Guangyu Zhao, Charlotte Wong, Shibo Jiang, Kwok-Yung Yuen, Dong-Yan Jin, and Bo-Jian Zheng. Cleavage of spike protein of sars coronavirus by protease factor xa is associated with viral infectivity. Biochemical and biophysical research communications, 359(1):174–179, 2007.

[11] Anthony R Fehr and Stanley Perlman. Coronaviruses: an overview of their replication and pathogenesis. In Coronaviruses, pages 1–23. Springer, 2015.

[12] Sayed S Sohrab and Esam I Azhar. Genetic diversity of mers-cov spike protein gene in saudi arabia. Journal of Infection and Public Health, 2019.

[13] Jianfei Chen, Xiaozhen Liu, D. Shi, Hongyan Shi, Xin Zhang, Changlong Li, Yanbin Chi, and Li Feng. Detection and molecular diversity of spike gene of porcine epidemic diarrhea virus in china. Viruses, 5(10):2601–2613, 2013.

[14] Beth N Licitra, Jean K Millet, Andrew D Regan, Brian S Hamilton, Vera D Rinaldi, Gerald E Duhamel, and Gary R Whittaker. Mutation in spike protein cleavage site and pathogenesis of feline coronavirus. Emerging infectious diseases, 19(7):1066, 2013.

[15] Xian-Dan Lin, Wen Wang, Zong-Yu Hao, Zhao-Xiao Wang, Wen-Ping Guo, Xiao-Qing Guan, Miao-Ruo Wang, Hong-Wei Wang, Run-Hong Zhou, Ming-Hui Li, et al. Extensive diversity of coronaviruses in bats from china. Virology, 507:1–10, 2017.

[16] Thanigaimalai Pillaiyar, Manoj Manickam, Vigneshwaran Namasivayam, Yoshio Hayashi, and Sang-Hun Jung. An overview of severe acute respiratory syndrome–coronavirus (sars-cov) 3cl protease inhibitors: Peptidomimetics and small molecule chemotherapy. Journal of medicinal chemistry, 59(14):6595–6628, 2016.

[17] Christopher Grow, Kaifu Gao, Duc Duy Nguyen, and Guo-Wei Wei. Generative network complex (gnc) for drug discovery. Communications in Information and Systems, 19:241–277, 2019.

[18] Duc Duy Nguyen, Kaifu Gao, Menglun Wang, and Guo-Wei Wei. Mathdl: Mathematical deep learning for d3r grand challenge 4. Journal of Computer-Aided Molecular Design, 2019, https://link.springer.com/article/10.1007/s10822-019-00237-5.

[19] Robin Winter, Floriane Montanari, Frank Noé, and Djork-Arné Clevert. Learning continuous and data-driven molecular descriptors by translating equivalent chemical representations. Chemical science, 10(6):1692–1701, 2019.

[20] David Rogers and Mathew Hahn. Extended-connectivity fingerprints. Journal of chemical information and modeling, 50(5):742–754, 2010.

[21] Joseph L Durant, Burton A Leland, Douglas R Henry, and James G Nourse. Reoptimization of mdl keys for use in drug discovery. Journal of chemical information and computer sciences, 42(6):1273–1280, 2002.

[22] Greg Landrum et al. Rdkit: Open-source cheminformatics, 2006.

[23] Duc Duy Nguyen, Zixuan Cang, Kedi Wu, Menglun Wang, Yin Cao, and Guo-Wei Wei. Mathematical deep learning for pose and binding affinity prediction and ranking in d3r grand challenges. Journal of computer-aided molecular design, 33(1):71–82, 2019.

[24] Duc D Nguyen, Zixuan Cang, and Guo-Wei Wei. A review of mathematical representations of biomolecular data. Physical Chemistry Chemical Physics, 2020, http://dx.doi.org/10.1039/C9CP06554G.

[25] Zhihai Liu, Yan Li, Li Han, Jie Li, Jie Liu, Zhixiong Zhao, Wei Nie, Yuchen Liu, and Renxiao Wang. Pdb-wide collection of binding data: current status of the pdbbind database. Bioinformatics, 31(3):405–412, 2014.

[26] Oleg Trott and Arthur J Olson. Autodock vina: improving the speed and accuracy of docking with a new scoring function, efficient optimization, and multithreading. Journal of computational chemistry, 31(2):455–461, 2010.

[27] Gareth Jones, Peter Willett, Robert C Glen, Andrew R Leach, and Robin Taylor. Development and validation of a genetic algorithm for flexible docking. Journal of molecular biology, 267(3):727–748, 1997.

[28] Richard A Friesner, Jay L Banks, Robert B Murphy, Thomas A Halgren, Jasna J Klicic, Daniel T Mainz, Matthew P Repasky, Eric H Knoll, Mee Shelley, Jason K Perry, et al. Glide: a new approach for rapid, accurate docking and scoring. 1. method and assessment of docking accuracy. Journal of medicinal chemistry, 47(7):1739–1749, 2004.

[29] Anna Gaulton, Louisa J Bellis, A Patricia Bento, Jon Chambers, Mark Davies, Anne Hersey, Yvonne Light, Shaun McGlinchey, David Michalovich, Bissan Al-Lazikani, et al. Chembl: a large-scale bioactivity database for drug discovery. Nucleic acids research, 40(D1):D1100–D1107, 2012.

[30] Li Di and Edward H Kerns. Biological assay challenges from compound solubility: strategies for bioassay optimization. Drug discovery today, 11(9-10):446–451, 2006.

[31] Jarmo Huuskonen, Marja Salo, and Jyrki Taskinen. Aqueous solubility prediction of drugs based on molecular topology and neural network modeling. Journal of chemical information and computer sciences, 38(3):450–456, 1998.

[32] William L Jorgensen and Erin M Duffy. Prediction of drug solubility from monte carlo simulations. Bioorganic & Medicinal Chemistry Letters, 10(11):1155–1158, 2000.

[33] Jacek Kujawski, Hanna Popielarska, Anna Myka, Beata Drabińska, and Marek K Bernard. The log p parameter as a molecular descriptor in the computer-aided drug design–an overview. Computational Methods in Science and Technology, 18(2):81–88, 2012.

[34] Christopher A Lipinski. Lead-and drug-like compounds: the rule-of-five revolution. Drug Discovery Today: Technologies, 1(4):337–341, 2004.

[35] Peter Ertl and Ansgar Schuffenhauer. Estimation of synthetic accessibility score of drug-like molecules based on molecular complexity and fragment contributions. Journal of cheminformatics, 1(1):8, 2009.

